# Midline Assembloids Reveal Regulators of Human Axon Guidance

**DOI:** 10.1101/2024.06.26.600229

**Authors:** Massimo M. Onesto, Neal D. Amin, Chenjie Pan, Xiaoyu Chen, Noah Reis, Alfredo M. Valencia, Zuzana Hudacova, James P. McQueen, Marc Tessier-Lavigne, Sergiu P. Paşca

**Affiliations:** Neurosciences Graduate Program, Stanford University, Stanford, CA 94305, USA; Department of Psychiatry and Behavioral Sciences, Stanford University School of Medicine, Stanford, CA 94305, USA; Stanford Brain Organogenesis, Wu Tsai Neurosciences Institute, Stanford University, Stanford, CA 94305, USA; Department of Biology, Stanford University, Stanford, CA 94305, USA

## Abstract

Organizers are specialized cell populations that orchestrate cell patterning and axon guidance in the developing nervous system. Although non-human models have led to fundamental discoveries about the organization of the nervous system midline by the floor plate, an experimental model of human floor plate would enable broader insights into regulation of human neurodevelopment and midline connectivity. Here, we have developed stem cell-derived organoids resembling human floor plate (hFpO) and assembled them with spinal cord organoids (hSpO) to generate midline assembloids (hMA). We demonstrate that hFpO promote Sonic hedgehog-dependent ventral patterning of human spinal progenitors and Netrin-dependent guidance of human commissural axons, paralleling non-human models. To investigate evolutionary-divergent midline regulators, we profiled the hFpO secretome and identified 27 evolutionarily divergent genes between human and mouse. Utilizing the hMA platform, we targeted these candidates in an arrayed CRISPR knockout screen and reveal that *GALNT2*, a gene involved in O-linked glycosylation, impairs floor plate-mediated guidance of commissural axons in humans. This novel platform extends prior axon guidance discoveries into human-specific neurobiology with implications for mechanisms of nervous system evolution and neurodevelopmental disorders.

## Introduction

Organizers serve as neurodevelopmental guideposts that regulate spatial patterning of neural tissues into distinct domains and the guidance of axons across long distances to their correct targets. The floor plate (FP) organizer, located ventrally along the neural tube midline, is comprised of specialized neuroepithelial cells that secrete morphogens and axon guidance cues into gradients along the dorsoventral axis. This property facilitates the patterning of the ventral spinal cord into distinct domains and guides axons across the midline to establish bilateral connectivity, which governs locomotive and sensory pathways.^1-4^

Although non-human models have provided fundamental insights into these developmental mechanisms, their organizer conservation and potential functional divergence in human midline development remains less understood.^5^ Human midline brain structures are particularly susceptible to decussation defects, for example agenesis of the corpus callosum or midline crossing defects during neurodevelopment.^6^ Therefore, developing models to directly study the human midline could advance our understanding of bilateral axon guidance and the molecular mechanisms that may contribute to neurodevelopmental disorders.

We previously developed an approach to model complex cellular interactions, including cell migration and axonal projections, during human brain development by generating regionalized neural organoids and subsequently integrating them *in vitro* to form three-dimensional (3D) assembloids.^7-13^ While previous studies have derived human FP cells, for instance as a precursor to generate midbrain dopamine neurons, their contribution to cell specification and axon guidance have been incompletely explored.^14^ In this study, we aimed to develop a protocol to generate functional human FP organoids (hFpO) and investigate their patterning and axon guidance effects on the developing human nervous system in assembloids.

We separately generated hFpO and spinal organoids (hSpO) and assembled them to form human midline assembloids (hMA). The integration of hFpO with early-stage hSpO induced sonic hedgehog (SHH)-dependent ventral patterning. Late-stage hSpO assembly with hFpO enabled recapitulation of the well-known guidance of commissural axons by Netrin-1 (NTN1), and also underscored a guidance role for F-spondin (SPON1), which has been reported as a cue for commissural axons. We comprehensively profiled the hFpO secretome via liquid chromatography mass spectrometry (LC-MS) and the hFpO transcriptome with single cell RNA sequencing (scRNA seq), identifying several evolutionarily divergent gene expression signatures enriched in the human floor plate. Using an arrayed CRISPR screen in hMA, we identified which of these factors regulate human FP-dependent axon guidance. Taken together, these experiments introduce a new human cellular model for investigating FP functions and its role in midline development. Understanding these mechanisms holds promise for elucidating both conserved and human-specific effectors governing midline axon guidance and the mechanisms establishing neural connectivity across species.

## Results

### Generation and characterization of hFpO and hSpO

To generate human organizer-like cells of the FP from human induced pluripotent stem (hiPS) cells, we exposed early neural organoids to a combination of growth factors and small molecules, mimicking neuralization and notochord-driven FP induction (Fig. 1a,b). We first enzymatically dissociated hiPS cells into single cells and aggregated ∼2,000 cells in ultra-low attachment round bottom 96-well plates. After initiating neuralization with the dual SMAD inhibitors dorsomorphin and SB-431542, we added 5 µM of smoothened agonist (SAG) to mimic secretion of SHH from the notochord, and supplemented with the growth factors FGF2, the WNT activator CHIR, and retinoic acid (Supplementary Fig. 1a). Immunocytochemistry of organoids at day 8 revealed a very high proportion (∼85%) of FOXA2^+^ nuclei (Fig. 1c,d). In parallel, we generated early-stage hSpO using an approach we previously developed^9^ with a protocol modification of removing any exogenous SHH activators to avoid the induction of ventral spinal progenitors (Fig. 1b, Supplementary Fig. 1a). RT-qPCR showed higher levels of the ventral transcription factors *FOXA2* and *NKX2-2*, the morphogen *SHH*, and the guidance molecule *NTN1* in hFpO than in hSpO (Fig. 1e). Moreover, hSpO showed no FOXA2 protein expression by immunohistochemistry and had higher expression of the dorsal transcription factors *PAX6* and *PAX3* by qPCR (Fig. 1d,e). We next verified the secretion of known FP morphogens and guidance molecules by incubating organoids in serum-free media for 24 hours and collecting the supernatant (conditioned media, CM). We performed western blot on organoid cell extracts and conditioned media collected from day 8 hFpO and hSpO and found that hFpO displayed enriched SHH expression and both cell-associated and secreted NTN1 (Fig. 1f). To assess the bioactivity of the secreted components of the hFpO on axon extension, we treated mouse dorsal spinal explants embedded in collagen gels with either hFpO or hSpO conditioned media for 40 hours. This assay revealed a pronounced increase in axon outgrowth from spinal explants treated with hFpO conditioned media compared to explants treated with conditioned media from hSpO (Fig. 1h,i).

**Figure 1:**
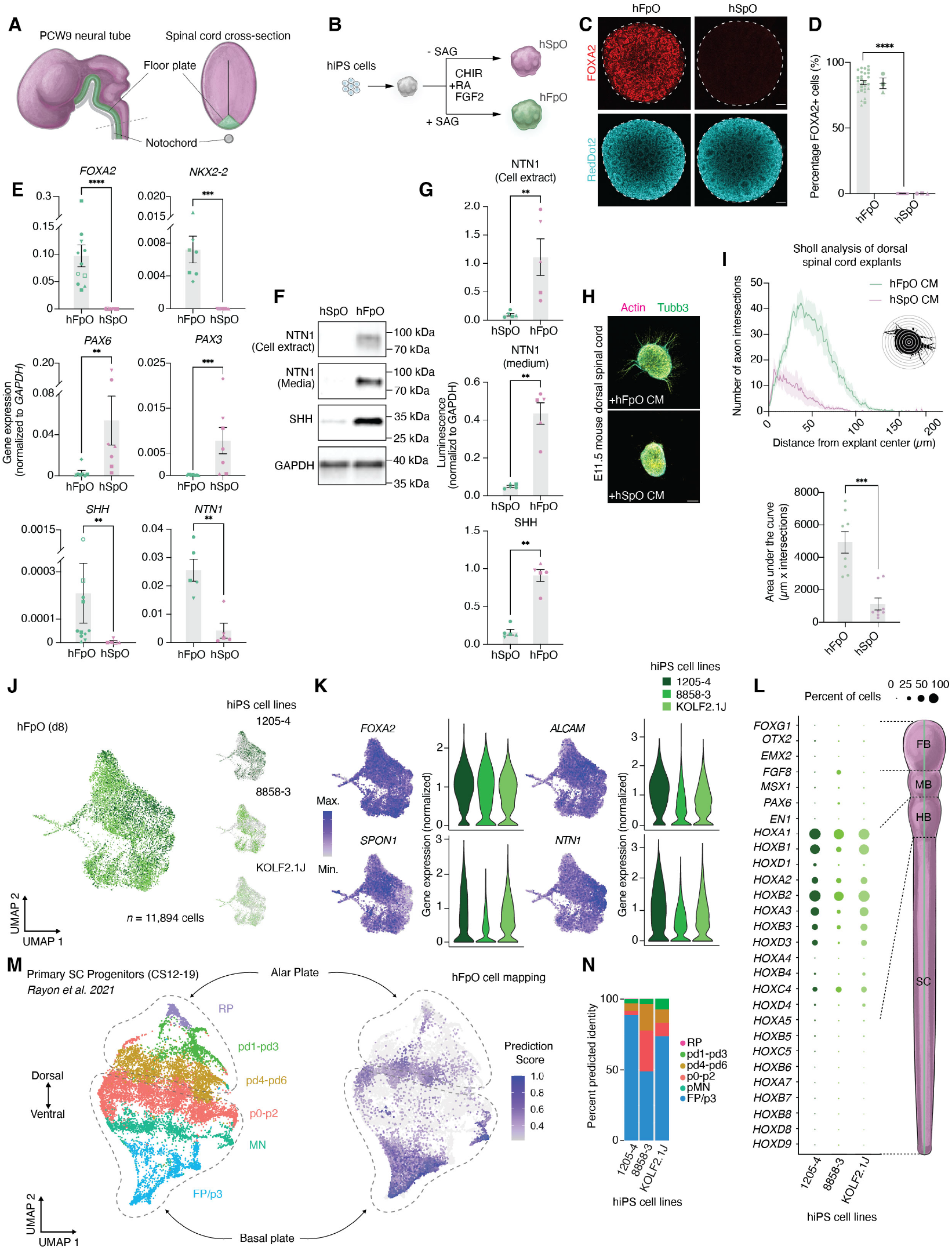
Generation and characterization of 3D hFpO and hSpO. **(A)** Schematic illustrating a developing human neural tube (purple) and location of the FP (green). **(B)** Schematic illustrating the generation of human FP organoids (hFpO, green) and human spinal organoids (hSpO, purple). **(C)** Representative immunocytochemistry images of FOXA2 expression and RedDot2 (nuclear stain) at day 8 of differentiation in cleared hFpO and hSpO. Scale bar, 100 μm. **(D)** Quantification of FOXA2 positive nuclei from immunocytochemistry images of individual hFpO and hSpO (n = 27 hFpO and 11 hSpO in 1-3 differentiation experiments from 3 hiPS cell lines; two-tailed t-test: ****P < 0.0001). Left bar shows individual organoids and the right bar indicates different hiPS cell line. Data is presented as mean ± SEM. **(E)** Gene expression analysis (by RT-qPCR) of *FOXA2, NKX2-2, PAX6, PAX3, SHH*, and *NTN1* in hFpO and hSpO at day 8 of in vitro differentiation. Two-tailed Mann-Whitney test was used. For *FOXA2*: n = 11 hFpO and 8 hSpO from 8 and 6 hiPS cell lines, respectively; ****P < 0.0001. For *NKX2-2*: n = 7 hFpO and 7 hSpO from 5 hiPS cell lines; ***P = 0.0006. For *PAX6*: n = 7 hFpO and 7 hSpO from 5 hiPS cell lines; **P = 0.0023. For *PAX3*: n = 7 hFpO and 7 hSpO from 5 hiPS cell lines; ***P = 0.0006. For *SHH*: n = 11 hFpO and 5 hSpO from 8 and 5 hiPS cell lines, respectively; **P = 0.0018. For *NTN1*: n = 5 hFpO and 5 hSpO from 4 hiPS cell lines; **P = 0.0019. Points indicate organoid samples from different differentiation experiments. Data is presented as mean ± SEM **(F)** Western blot of SHH and NTN1(cell extract and conditioned media) collected from hFpO and hSpO. Immunoblotting of GAPDH was used as loading control. **(G)** Western blot quantification of SHH and NTN1 (cell extract and conditioned media) from hFpO and hSpO normalized to GAPDH luminescence. Data is presented as mean ± SEM. Two-tailed Mann-Whitney test was used. For NTN1 (cell extract): n = 5 hFpO and 5 hSpO from 5 hiPS cell lines; **P = 0.0079. For NTN1 (media): n = 5 hFpO and 5 hSpO from 5 hiPS cell lines; **P = 0.0079. For SHH: n = 5 hFpO and 5 hSpO from 5 hiPS cell lines; **P = 0.0079. **(H)** Representative z-projected confocal images of E11.5 mouse dorsal spinal explants cultured for 40hr in conditioned media (CM) collected from either hFpO or hSpO. Explants were stained for Tubb3 and phalloidin (actin stain) to visualize axons. Scale bar, 100 μm. **(I)** Sholl analysis of axon outgrowth from conditioned media treated E11.5 mouse dorsal spinal explants and area under the curve quantification. Two-tailed Mann-Whitney test ***P = 0.0003; n = 8 for hSpO and n = 8 hFpO CM treated explants dissected from 2-3 embryos. **(J)** UMAP visualization of single cell gene expression of hFpO at day 8, separately colored by individual cell lines (shades of green). (n = 11,894 cells from 3 hiPS cell lines) **(K)** UMAP and violin plots showing gene expression of selected FP markers across 3 hiPS cell lines (shades of green). Color scale indicates normalized gene expression level. **(L)** Dot plots showing the expression of rostral-caudal defining genes expressed in hFpO cells. The size of the circle represents the percent of cells expressing each gene in 3 hiPS cell lines (shades of green). **(M)** UMAP visualization of subsetted primary human spinal progenitor clusters (left, CS12-19) and mapped hFpO cells (day 8) onto the primary atlas (right, color scale indicates prediction score of mapped hFpO cells and light gray points indicates reference atlas cells). **(N)** Bar plot displaying the proportion of predicted domains of hFpO cells from mapping onto a primary spinal progenitor atlas in each of the three hiPS cell lines.

To study cell diversity in hFpO, we performed droplet-based single cell RNA sequencing (scRNA-seq) analysis at day 8 of differentiation (n = 11,894 cells, 3 hiPS cell lines; Fig. 1j). We found that cells were generally homogenous in their gene expression signature, with high expression of known FP markers *FOXA2, ALCAM, SPON1*, and *NTN1* (Fig. 1k) resembling the caudal FP (Fig. 1l). We next mapped the hFpO cells onto a primary fetal atlas spanning CS12 to CS19 of the human spinal cord and found high overlap (up to 90%) with the ventral domain cluster (FP/p3; Fig. 1m,n, Supplementary Fig. 1b). Additionally, we found that hFpO cells resemble the expression signatures of human fetal FP cells at stage CS12 (Supplementary Fig. 1c). We also used VoxHunt^15^ and found that the hFpO cells preferentially mapped to the ventral portion of the E11 mouse hindbrain where the FP resides (Supplementary Fig. 1d). Taken together, these experiments suggest that we can reliably and efficiently generate 3D cultures resembling the early-stage human FP from stem cells.

### Generation of hMA to model patterning

To model the influence of the FP on ventral spinal cord patterning, we created assembloids using hFpO derived from a fluorescent hiPS cell line and hSpO from a non-fluorescent line of the same background. We placed a day 8 hFpO and a day 8 hSpO in proximity in a low-attachment round-bottom well, where they assembled within 24 hours (Fig. 2a). The day 8 timepoint for the hSpO assembly was chosen because it composed primarily of unpatterned neural progenitors, ensuring that any influence on ventral induction would result from the effects of the hFpO rather than from exogenous molecules added to the medium. We collected assembloids at 3 and 7 days after fusion (d.a.f.) and assessed ventral progenitor induction on the non-fluorescent hSpO side using immunocytochemistry. We observed a gradient expression of the ventral transcription factors *FOXA2, NKX2*.*2*, and *NKX6*.*1* at the hFpO-hSpO boundary suggestive of FP-induced ventral patterning (Fig.2b,c). To verify this, we exposed assembloids to the Smoothened inhibitor (SHH-pathway inhibitor), cyclopamine, during fusion and found that it completely abolished expression of ventral markers in hSpO fused to hFpO (Fig. 2b,c). Moreover, fusion of a fluorescent hSpO with a non-fluorescent hSpO had no effect, further confirming the specific ventral inductive effect of hFpO (Supplementary Fig. 2a). To investigate hFpO-mediated ventral patterning at the transcriptomic level, we sorted cells for scRNA-seq (Fig. 2d, Supplementary Fig. 6a-c). hSpO sample clusters were distinct: the unassembled hSpO primarily expressed dorsal neural progenitor transcription factors (*LMX1A, PAX7, PAX3, IRX3, PAX6*), while the assembled hSpO primarily expressed ventral progenitor transcription factors (*NKX6*.*2, OLIG2, NKX2*.*2, FOXA2;* Fig. 2e-g). We also examined *GLI1* and *GLI3* expression profiles as proxies for SHH-pathway activation and inhibition, respectively. We discovered a positive correlation between ventral transcription factors and *GLI1* expression, and a negative correlation with *GLI3* expression within hSpO. Dorsal transcription factors showed the opposite correlation, indicating an inverse relationship between *GLI1/GLI3* and ventral/dorsal transcription factor expression (Fig. 2h). We next mapped hSpO cell clusters onto a primary spinal cord atlas^16^ and found that the unassembled hSpO mainly resembled cells from dorsal spinal cord progenitor domains, while the hSpO in the assembloid mostly resembled ventral-mapping progenitors (Fig. 2i,j; Supplementary Fig. 2b). Gene expression profiling of the mapped assembled hSpO cells showed a high correlation with their respective domains of the primary spinal cord (Fig. 2k,l; Supplementary Fig. 2c). Overall, these experiments demonstrate that hFpO has organizer-like properties, inducing ventral spinal cord fates in assembloids. fluorescent hSpO cells assembled with non-fluorescent hFpO and stage-matched unassembled fluorescent hSpO

**Figure 2:**
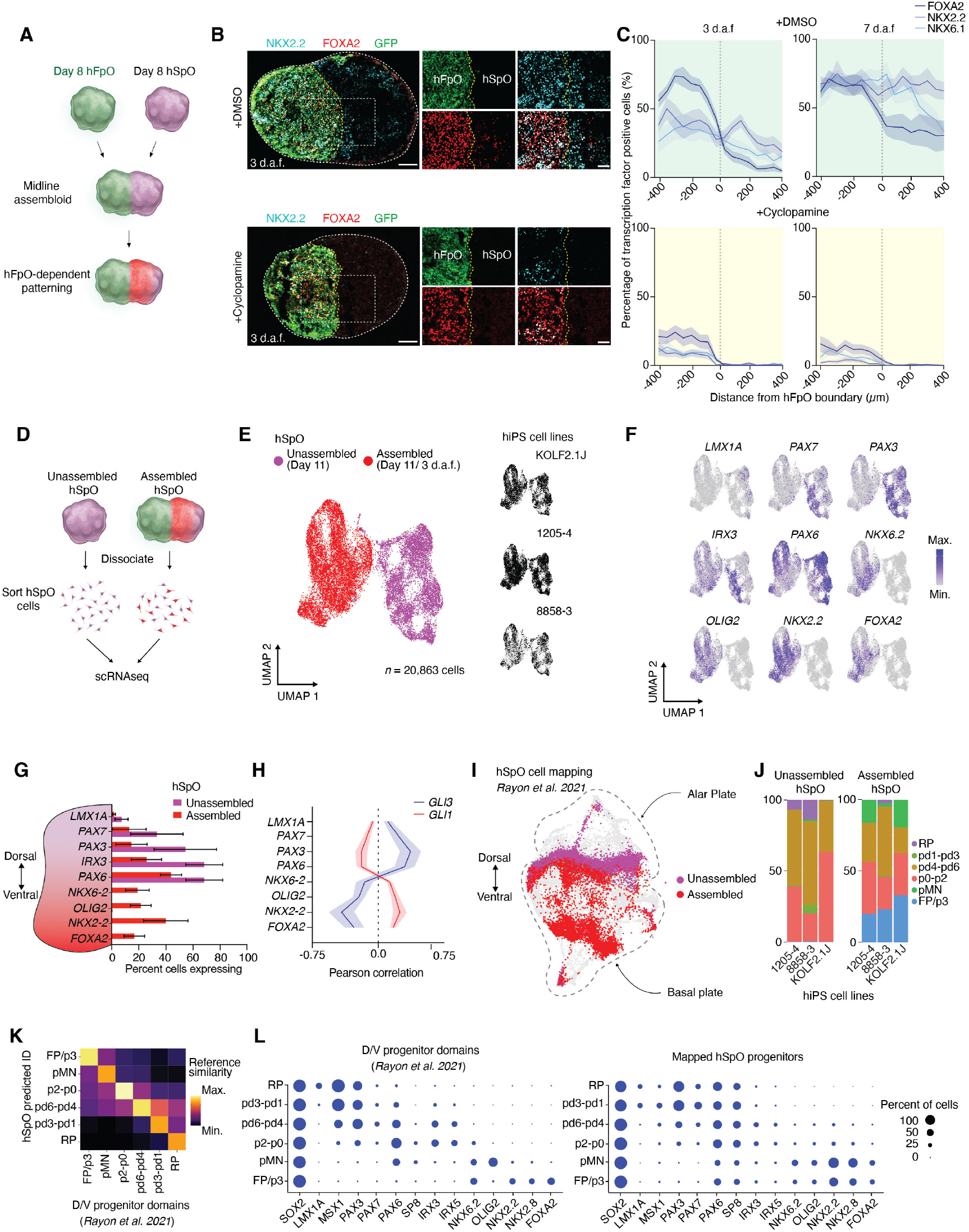
Generation and characterization of hMA for assessing SHH-dependent ventral patterning. **(A)** Schematic illustrating the assembly of day 8 hFpO (green) and hSpO (purple) to form hMA with hFpO-dependent patterning (red). **(B)** Representative widefield images of sectioned hMA cultured with and without cyclopamine at 3 days after fusion (3 d.a.f.). Sections were immunostained for NKX2.2 and FOXA2 and an endogenously fluorescent hiPS cell line was used to label the hFpO. Scale bar, 200 μm. **(C)** Plot profiles of the percent positive cells for ventral transcription factors (TF) from the hFpO-hSpO boundary (dotted line) 3 to 7 d.a.f in hMA cultured in the presence of DMSO or cyclopamine. Data is presented as mean ± SEM. For FOXA2 (dark blue), NKX2.2 (medium blue), and NKX6.1 (light blue): n = 7-9 assembloids per condition from 3 differentiation experiments from 1 hiPS cell line. **(D)** Schematic illustrating the process of collecting either unassembled hSpO or hFpO-assembled hSpO cells for scRNA seq analysis. **(E)** UMAP visualization of single cell gene expression of cells collected from unassembled (purple) and hFpO-assembled hSpO (red). (n = 20,863 cells from 3 hiPS cell lines). **(F)** UMAP visualization of selected dorsal and ventral spinal progenitor gene expression in unassembled and hFpO-assembled hSpO. Color scale indicated normalized gene expression level. **(G)** Bar plot displaying the percentage of cells expressing selected dorsal and ventral spinal progenitor genes in unassembled (purple bar; n = 3 samples) and hFpO-assembled hSpO (red bar; n = 3 samples) from 3 hiPS lines. Data is presented as mean ± SEM. **(H)** Pearson correlation plot of selected dorsal and ventral spinal progenitor gene expression with *GL1* (blue line) or *GLI3* (red line) expression in unassembled (n = 3 samples) and hFpO-assembled hSpO (n = 3 samples) across 3 hiPS cell lines. Data is presented as mean ± SEM. Two-way ANOVA and Bonferroni’s test for multiple comparison, F_1,32_ = 0.5113, P = 0.4789. For *LMX1A*, P > 0.9999. For *PAX7*, *P = 0.0236. For *PAX3, \*\**\*P = 0.0002. For *PAX6*, ***P = 0.0010. For *NKX6*.*2*, P = 0.1313. For *OLIG2*, **P = 0.0065. For *NKX2*.*2*, ****P < 0.0001. For *FOXA2*, **P = 0.0029. **(I)** UMAP visualization of unassembled hSpO (purple) and hFpO-assembled hSpO (red) onto a primary human spinal progenitor atlas (light gray). **(J)** Bar plots of the proportion of unassembled (n = 3 samples) and hFpO-assembled (n = 3 samples) hSpO predicted progenitor cell domain identity from a human spinal progenitor atlas from 3 hiPS cell lines. **(K)** Heatmap depicting the correlation of gene expression profiles between predicted hSpO progenitor domains with primary human spinal cord progenitor domains. **(L)** Dot plots comparing the expression of dorsal-ventral defining genes of mapped unassembled and assembled hSpO cells (right) vs primary human spinal cord progenitor domains (left). The size of the circle represents the percent of cells expressing each gene in 3 hiPS cell lines.

### Modeling axon guidance in hMA

To model axon guidance in hMA, we first verified the presence key populations of neurons that project contralaterally in the spinal cord in hSpO. This required using hSpO at a later stage while excluding SHH-agonists to generate cell populations that resemble dorsal spinal cord interneurons (Supplementary Fig. 3a). Single cell profiling of hSpO at day 35 revealed clusters of cells expressing specific markers of dI1, dI2, dI3, dI4, and dI5 interneuron populations (Fig. 3b,c). One cluster expressed markers of contralateral projecting dorsal interneurons known in mouse to project axons to the FP, including *ROBO3, TAG-1, EBF2*, and *NHLH2*, (Fig 3b,d; Supplementary Fig. 3b,c).^17-20^ Immunocytochemistry for ROBO3 expression revealed a temporally-regulated expression pattern in hSpO, which is consistent with prior reports from mouse studies.^21^ In contrast, expression of the neuronal marker TUBB3 showed a steady increase across this period (Fig. 3e,f). To assess whether hiPS cell-derived commissural neurons project from hSpO to the hFpO, we assembled hSpO when ROBO3 expression is approaching its peak at day 22, with day 8 hFpO (Fig. 3a,g). Remarkably, we observed progressive and directed axonal projections and fasciculation of axon bundles invading the hFpO (Fig. 3g,h). To determine if these projections depended on hFpO, we assembled hSpO at day 22 with day 8 hSpO which lacks FP cells and found only rare projecting commissural axons with a defasciculated appearance (Fig. 3g,h). Quantification of hSpO axon projection directionality within the two preparations revealed much higher levels of ROBO3^+^ projections directed towards the hFpO (Fig. 3i), indicating preferential guidance of hSpO commissural neurons in an hFpO-dependent manner (Fig. 3i).

**Figure 3:**
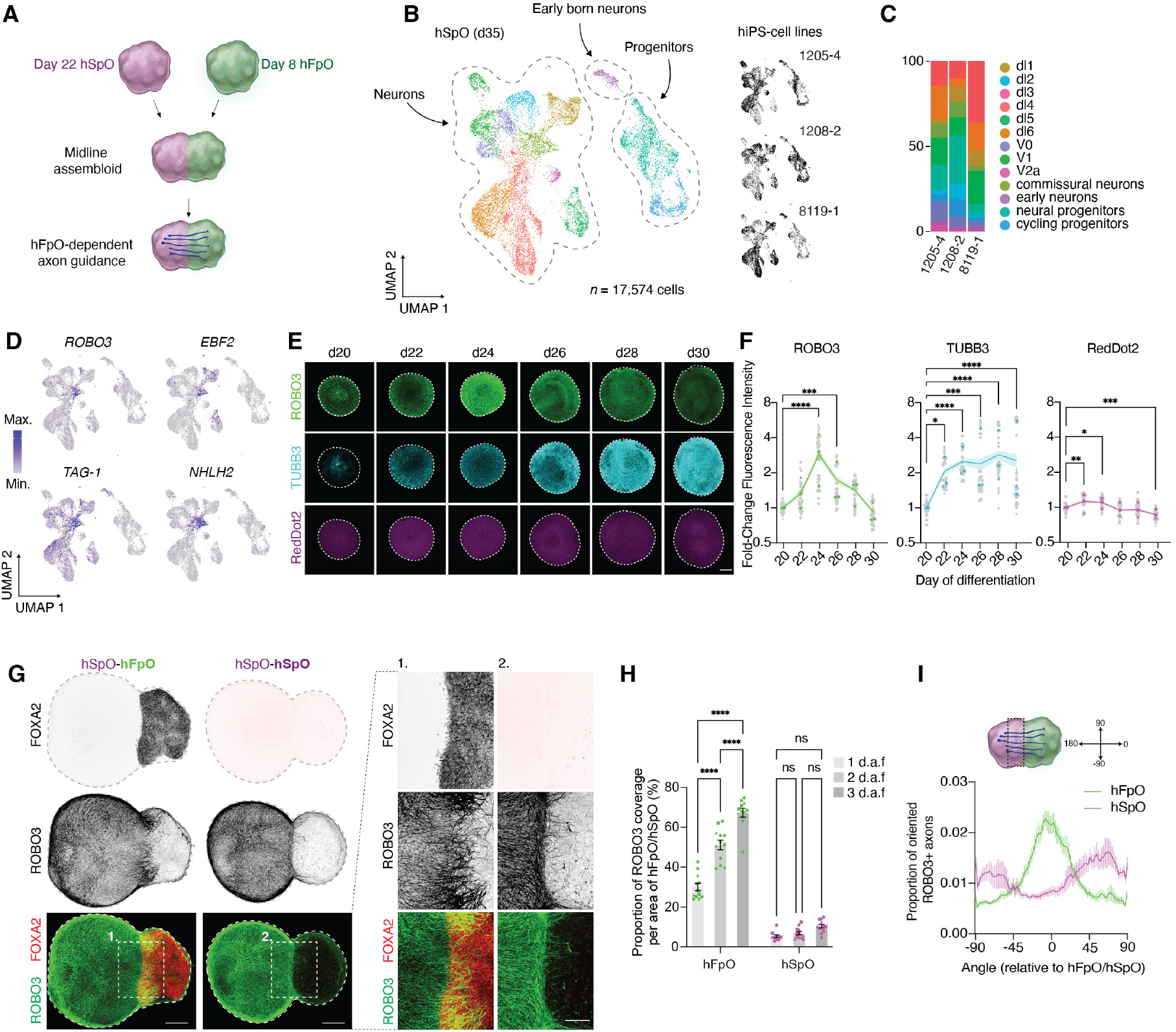
Generation and characterization of hMA for assessing hFP-dependent axon guidance. **(A)** Schematic illustrating the assembly of day 8 hFpO (green) and day 22 hSpO (purple) to form hMA for modeling hFpO-dependent axon guidance. **(B)** UMAP visualization of single cell gene expression of cells collected from day 35 (d35) hSpO. (n = 17,574 cells from 3 hiPS cell lines). **(C)** Bar plots of the proportion of annotated d35 hSpO cell clusters across 3 hiPS cell lines. **(D)** UMAP visualization of single cell gene expression of markers enriched in dorsal commissural neuron populations, *ROBO3, EBF2, TAG-1*, and *NHLH2*. Color scale indicates normalized gene expression level. **(E)** Representative z-projected immunocytochemistry images of cleared hSpO showing expression of ROBO3 and TUBB3 with RedDot2 nuclear stain from day 20 to day 30 of differentiation. Scale bar, 200 μm. **(F)** Quantification of fold-change in ROBO3, TUBB3 and Reddot2 fluorescence relative to day 20 of differentiation measured from 22–32 individual organoids from 3 hiPS cell lines (individual samples shown as gray points, mean per line is shown as a colored points, and the dashed line shows mean and SEM across samples). Line indicates mean while shaded area represents SEM. Plotted values are Log2 transformed. One-way ANOVA followed by Dunnett’s multiple comparison test was used. For ROBO3: F_5,161_ = 39.26, P <0.0001; ***P = 0.0003, ****P < 0.0001. For TUBB3: F_5,161_ = 6.864, P < 0.0001; *P = 0.0164, ***P = 0.0005, ****P <0.0001. For Reddot2: F_5,161_ = 17.81, P < 0.0001; **P = 0.0036, *P = 0.0171, ***P = 0.0008. Scale bar, 200 μm. **(G)** Representative z-projected immunocytochemistry images of ROBO3 and FOXA2 expression in cleared hMA expression at 2 days after fusion (d.a.f.). Scale bar, 200 μm; inset, 100 μm). **(H)** Quantification on the proportion of ROBO3+ axon coverage normalized to the area of either the assembled hFpO (green) or hSpO (purple) over time. Data points represent individual assembloids (n = 11-12 per condition from 2 hiPS cell lines) while bar plots indicate the mean at 1 d.a.f. (light gray), 2 d.a.f. (medium gray) and 3 d.a.f. (dark gray). Comparison of the effects across time points in each condition; Two-way ANOVA and Tukey’s test for multiple comparison, F_2,63_ = 79.23, P < 0.0001; ****P < 0.0001. **(I)** Quantification of the amount of oriented ROBO3^+^ axons from the hSpO relative to either an assembled hSpO or hFpO at 2 d.a.f. (n = 12 per condition from 2 hiPS cell lines). Zero degrees indicates projection directly towards and +-90 degrees indicates away from the assembled hFpO/hSpO. The analyzed ROI is indicated by the dotted line on the hMA schematic. Two-tailed Mann-Whitney test comparing means at 0 degrees: ****P < 0.0001.

### Identification of human enriched FP genes

We leveraged the ability of our hFpO system to generate massive numbers of highly pure FP cells at scale for a proteomic analysis of membrane and secreted proteins enriched in the human FP compared to mouse. We collected CM from hFpO from three hiPS cell lines (about 30 million hFpO cells) and performed LC-MS which captured 1,876 proteins (Supplementary Table 1). To identify which of these proteins are selectively enriched in FP cells, we identified genes transcriptomically detected in scRNA-seq of primary human FP. We retained genes that were enriched in FP versus other spinal progenitors and that were present in at least 30% of primary FP cells, resulting in a list of 415 FP-enriched genes (Supplementary Table 1). Of these, 326 encoded membrane and secreted proteins (Supplementary Fig. 4a) and 98 are known to be genes essential for cell survival (Depmap Portal, Supplementary Table 1). To determine if any of these FP-enriched genes are specific to humans, we first analyzed the conservation and divergence of gene expression patterns in human versus mouse FP cells^22^ and found that 8.6% of genes displayed a greater than 10-fold divergence in expression (Fig. 4a, Supplementary Fig. 4b, Supplementary Table 2). Interestingly, 3 of these genes had human specific expression in the human FP (*EFEMP1, TTR, CLSTN3)* (Fig. 4a, Supplementary Fig. 4c). Overall, these proteomic and transcriptomic experiments identified genes encoding membrane and secreted proteins that are enriched or specific to the human FP compared to the mouse FP.

**Figure 4:**
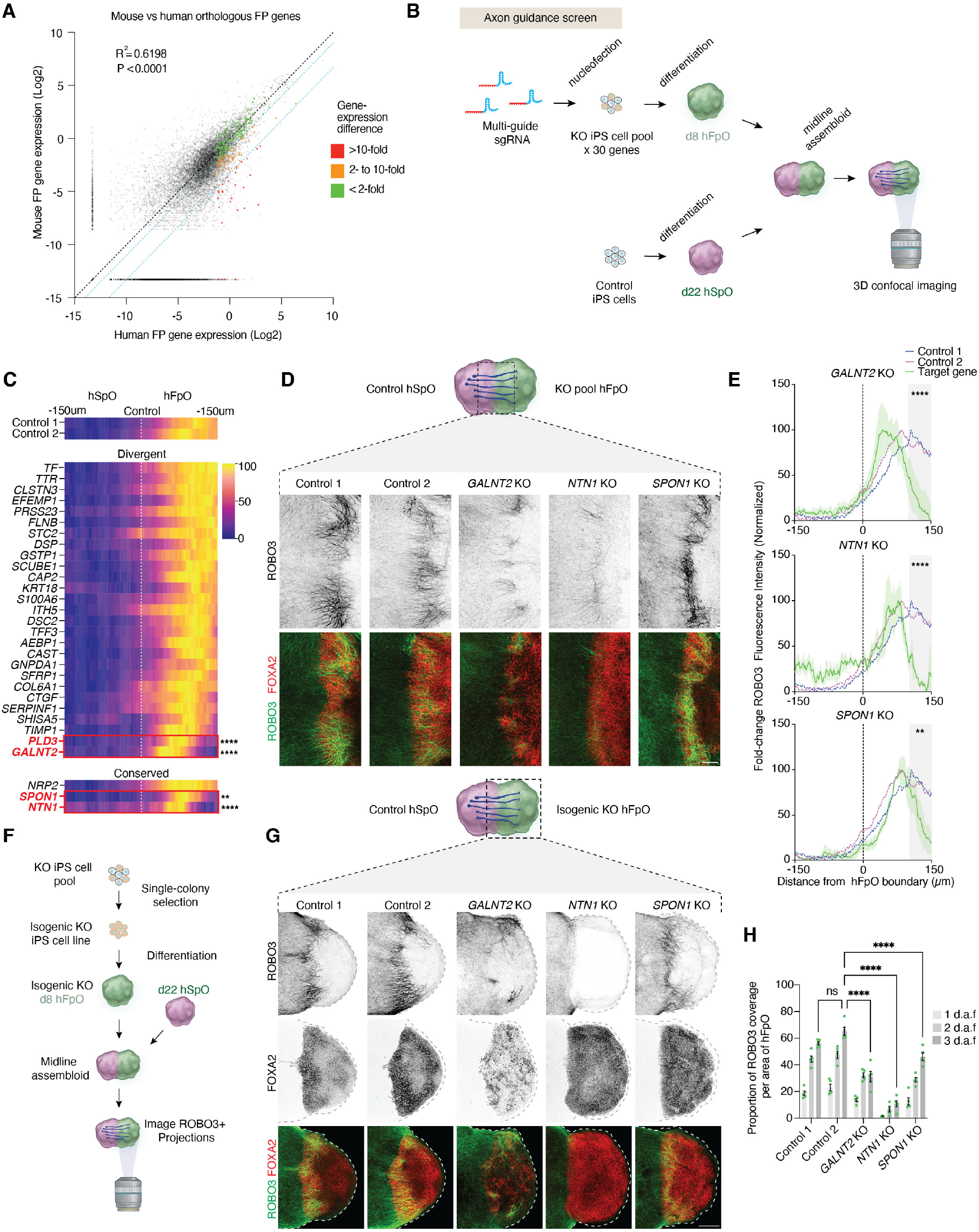
Screening human-enriched genes as effectors of hFP-mediated axon guidance. **(A)** Comparison of gene expression levels of 14,852 orthologous genes (gray points) between mouse (E9-14) and human (CS12-19) FP cells. Black dotted line indicates 1:1 gene expression while the blue dotted lines indicate 2-and 10-fold human enrichment. Colored points indicate membrane and secreted FP-enriched genes and indicate degree of gene expression divergence between human and mouse. **(B)** Schematic of workflow for generating hFpO from pooled KO hiPS cells and subsequently assembling with day 22 hSpO to form hMA in an arrayed format. At 3 d.a.f., the assembloids were cleared and confocal imaged to visualize commissural axons guiding towards the hFpO. **(C)** Heatmap displaying min/max normalized, fold-change in ROBO3 fluorescence intensity of hSpO axons 150 μm pre- and 150 μm post-crossing into the hFpO (n = 2-3 assembloids per gene). Two-way ANOVA performed on measurements the most distal 50 μm of heatmap; F_31,1972_ = 10.05, P < 0.0001, following Dunnett’s multiple comparison test comparing Control 1 to all other conditions: ****P <0.0001 for Control 1 vs *GALNT2*, ****P <0.0001 for Control 1 vs *PLD3*, ****P < 0.0001 for Control 1 vs *NTN1*, and **P = 0.0025 for Control 1 vs *SPON1*. **(D)** Representative z-projected immunocytochemistry images of control and pooled KO hMA expressing ROBO3 and FOXA2 to visualize ROBO3+ axons projecting across the hFpO boundary. Scale bar, 100 μm. **(E)** Plot profiles of normalized fold-change ROBO3 fluorescence intensity of KO pool hFpO hMA 150 μm pre- and 150 μm post-crossing into the hFpO from (C). Dotted lines indicate the non-targeting control groups while the solid lines indicate the target gene group. Data is presented as mean ± SEM. Gray shaded region indicates the 50 μm bin where ROBO3 expression as a measure of axon guidance was assessed for statistical significance in (C). **(F)** Schematic outlining the method for generating isogenic KO hiPS cell lines from the axon guidance screen candidates and subsequent isogenic validation of phenotypes identified in the axon guidance screen. **(G)** Representative z-projected immunocytochemistry images of control and isogenic KO hMA expressing ROBO3 and FOXA2 at 2 d.a.f. Scale bar, 200 μm. **(H)** Bar plot displaying the proportion of ROBO3 coverage relative to the area of the isogenic KO hFpO candidates from 1-3 d.a.f (shades of grey; n = 4-5 assembloids per condition). Data is presented as mean ± SEM. Two-way ANOVA, F_4,59_ = 121.8, P < 0.0001, following Tukey’s multiple comparison test ****P < 0.0001.

### Arrayed CRISPR screen of membrane and secreted genes reveals regulators of human FP-dependent axon guidance

To investigate the role of human-enriched membrane and secreted genes in commissural axon guidance to the FP, we designed an arrayed CRISPR screen leveraging the accessibility of hMA for genetic manipulation and multiple functional readouts of FP-dependent axon guidance (Fig. 4b). We designed guides targeting 27 human FP genes with the highest divergence in gene expression compared to mouse FP and three conserved genes (*NTN1, SPON1*, and *NRP2*) previously implicated in FP-dependent axon guidance^23-25^ as well as non-targeting control (NTC) guides (Supplementary Table 3). We individually screened hiPS cells knockout pools for editing efficiency by inference of CRISPR edits (ICE) analysis and KO-score and found 90% exhibited an editing efficiency of >75% (Supplementary Fig.5a).^26^ We generated hFpO organoids from individual hiPS cell KO pools, and at day 8 of differentiation, we assembled them with unedited day 22 hSpO (Fig. 4b). Three days later, we fixed hFpO-hSpO assembloids, performed whole mount assembloid clearing, and immunostained for ROBO3 and FOXA2 to assess commissural axon projections (Fig. 4b). We imaged entire intact assembloids and measured fluorescence intensity of ROBO3 within 150 µm of the FOXA2 boundary, normalizing to pre-hFpO ROBO3+ fluorescence intensity (Fig. 4c-e). As expected, loss of *NTN1* and *SPON1* resulted in decreased ROBO3+ axon projection into the hFpO, consistent with their effects on commissural axon guidance in non-human model organisms (Fig. 4c-e).^18,23,24^ In contrast to a report a from rodent models loss of *NRP2* had no detectable effect on human commissural axon guidance in our preparation (Fig. 4c-e).^25^

An analysis of all targets identified two human-enriched FP gene candidates that strongly affected commissural axon projection: *GALNT2* and *PLD3* (Fig. 4c). *GALNT2* encodes an enzyme that initiates O-linked glycosylation by transferring N-acetylgalactosamine to proteins, and *PLD3* encodes a protein involved in lysosomal function and catalyzes the hydrolysis of phospholipids. *GALNT2* knockout reduced *FOXA2* gene expression at the transcript and protein levels, suggesting an important role in FP cell specification (Supplementary Fig. 5b-d). To validate these findings, we generated isogenic *GALNT2, NTN1*, and *SPON1* homozygous knockout hiPS cell lines and reproduced these axon projection phenotypes in hMA (Fig. 4f). Isogenic knockout was confirmed by ICE analysis as well as Western blot for target genes in cells and conditioned media from hFpO (Supplementary Fig. 5e,f). The isogenic knockouts showed significant reduction in commissural axon projection to the hFpO (Fig. 4g,h). Moreover, *GALNT2* knockout resulted in reduced *FOXA2* and *NKX2*.2 expression, along with a trend toward increased *PAX6* expression, indicating deficits in ventral neural patterning in hFpO (Supplementary Fig.5g). Together, these results suggest that hMA can be used to model human ventral neural patterning and axon guidance and to discover functionally important genes evolutionarily enriched in the human FP.

## Discussion

Stem cell-derived neural organoids are capable of recapitulating human cellular diversity across regions of the developing nervous system. The integration of multiple organoids within neural assembloids highlights the ability to capture aspects of self-organization *in vitro* by modeling intricate cell-cell interactions essential for brain connectivity and function. Our previous work has showcased this potential by modeling interneuron cell migration in forebrain assembloids and by modeling long-range projections and circuit formation in cortico-striatal, cortico-motor, cortico-thalamic, and ascending sensory pathway assembloids.^7-13^ However, the use of assembloids to explore patterning by organizer-like cells and the guidance of axons at the midline has not been explored.

Here, we developed stem cell-derived human hMA that model guidance of commissural axons to the FP, an essential step for left-right neural connectivity in bilateral nervous systems. First, we generated human FP organoids and defined their transcriptome and secretome profiles across multiple hiPS cell lines. Secondly, we developed a novel assembloid model that offers flexibility in the timing of integration to capture different aspects of midline development. By integrating hFpO with early-stage, unpatterned hSpO, we modeled SHH-dependent ventralization of spinal progenitors. Earlier studies employing hiPS cells to investigate ventral patterning predominantly relied on ectopic overexpression of a single recombinant morphogen gene, without taking into account the full compendium of secreted organizing factors that govern ventral midline development.^27,28^ By integrating hFpO with dorsal hSpO containing ROBO3-positive commissural neurons we modeled midline axon guidance towards the FP, recapitulating the guidance contributions of NTN1 and SPON1. Thirdly, we applied our hMA model to identify genes that control midline axon guidance specifically in human. We identified 27 evolutionarily divergent genes enriched in the human FP compared to mice and conducted an arrayed CRISPR knockout screen. This revealed an important role for the glycosylation enzyme GALNT2 in FP generation and commissural axon guidance to the FP. This may be mediated by GALNT2-dependent glycosylation of components of the SHH-pathway, a mechanism that has been suggested in the paralogous protein GALNT1.^29^ The ability of GALNT2 to regulate glycosylation of axon guidance molecules such as NTN1 has not been explored but could be an important modifier of midline connectivity as well.^30^ These findings underscore the utility of assembloids for arrayed CRISPR screens not only to probe axon guidance effectors but also suggests evolutionarily divergent gene expression that contribute to species-specific differences in midline formation and connectivity.

The human cellular model presented here holds potential for additional applications. The hFpO as well as hMA can be used to investigate differences in secreted versus cell-contact dependent mechanisms in ventral patterning that impact short versus long range signaling capabilities in ventral patterning^31^ and axon guidance.^32,33^ Furthermore, this model could allow the investigation of growth cone response switches at intermediate targets.^4^ Moving forward, it will be important to explore whether hFpO can recapitulate aspects of circuit assembly following axonal crossing. Lastly, comparison of primate and non-primate hFpO could elucidate the conserved and divergent molecular mechanisms of midline development and how they may contribute to midline defects frequently observed in humans by neuroimaging and postmortem histological analyses.

There are several limitations of this model. The current assembloid does not capture post-crossing axonal events, such as rostrocaudal turning and synaptogenesis. This could be achieved by generating a three-part assembloid that positions the hFpO in between two hSpO. Such a system could be used to study both switching of receptors as well as early contralateral connectivity. Studies of later hFpO stages within assembloids are limited by the need to move cultures to a neurogenesis-promoting media which may deplete FP populations over time since they have the capacity to function as neural precursors. Future studies can optimize the components of hMA media to promote neurogenesis of commissural neurons in the hSpO without interfering with hFpO cell fate. This will allow for longer culture of hMA and facilitate the study of human midline development and circuit formation.

Expanding beyond the spinal cord midline, the utilization of organizer regions in assembloids can be extended to model other brain structures. Notably, there is a gap in our understanding of organizers modeling the corpus callosum, the largest axonal tract in the human nervous system. While attempts have been made to emulate corpus callosum formation they lack the complex signaling events that govern axon guidance across the glial wedge.^34^ Furthermore, organizers that model the roof plate and the isthmic organizer could be used to study dorsal and anterior-posterior patterning, respectively. Ultimately, organizer organoids offer a versatile toolset to advance our knowledge of human cell specification, axon guidance, and evolution, which are crucial to understand human neurodevelopment and the impact of neurodevelopmental disorders.

## Acknowledgements

This work was supported by NINDS K08-NS123544-01 (N.D.A), Brain and Behavior Research Foundation (N.D.A), Foundations of the National Institutes of Health Deeda Blair Research Initiative (N.D.A.), the Stanford Brain Organogenesis Program in the Wu Tsai Neuroscience Institute and Bio-X (S.P.P), Kwan Funds (S.P.P), Senkut Funds (S.P.P), the Ludwig Foundation (S.P.P), the Mann Foundation (S.P.P), the New York Stem Cell Foundation (NYSCF) (S.P.P), the Stanford Science Fellows Program (A.M.V.), and Ford Foundation Postdoctoral Fellowship (A.M.V.). S.P.P. is Chan Zuckerberg Initiative (CZI) Ben Barres Investigator and a CZ BioHub Investigator. This paper was typeset with the bioRxiv word template by @Chrelli: www.github.com/chrelli/bioRxiv-word-template

## Author contributions

M.O., N.A., M.T.L., and S.P.P. conceived the project and designed the experiments. M.O. and N.A. developed the hFpO differentiation and characterized these cultures. M.O. conducted all differentiation, sample collection, mouse embryo dissection/culture, and scRNA seq analysis. M.O. analyzed the evolutionarily divergent FP genes and generated the list of genes used in the screen. M.O. edited all CRISPR KO hiPSC lines (with support from A.V. and X.C.), generated hMA and performed clearing/imaging of all samples during the screen. M.O., A.V., and Z.H. performed genotyping of hFpO cell pools during the screen. M.O. and X.C. performed FACS of hMA. N.R. and J.P.M. performed RT-qPCR for hFpO/hSpO. N.R. performed imaging analysis for the screen. C.P. performed western blotting, maintained mouse colonies, and provided timed mouse embryos. M.O. and S.P.P. wrote the manuscript with input from all authors. S.P.P. supervised all aspects of this study.

## Materials and Methods

### Culture of hiPS cells

The hiPS cell lines utilized in this study were validated following previously established methods. The 2242-1, 1205-4, 8858-3, 0524-1, 0307-1, 8119-1, 1208-2 control hiPS cell lines were used to generate hFpO and/or hSpO, and were derived at Stanford University. The KOLF2.1J control stem cell line was used to generate hFpO and/or hSpO and was obtained from Jackson Laboratory (Farmington, Connecticut). The CAG::GFP cell line was engineered from the parental cell line 8858-3.

In brief, hiPS cells were cultured in six-well plates coated with recombinant human vitronectin (Life Technologies, catalogue no. A14700) using Essential 8 medium (Life Technologies, catalogue no. A1517001). For passaging hiPS cell colonies, cells were treated with 0.5 mM EDTA for 7 min at room temperature, resuspended in Essential 8 medium, and transferred to new six-well plates. Regular testing and maintenance ensured the cultures remained free of mycoplasma. Approval for the derivation and use of these cell lines was obtained through the Stanford Institutional Review Board. Validation of hiPS cell genome integrity was conducted using high-density single-nucleotide polymorphism arrays.

### Generation of hFpO from hiPS cells

hFpO were differentiated from feeder-free maintained hiPS cells, as previously described. To generate 3D hFpO, hiPS cells were dissociated with Accutase (Innovative Cell Technologies, AT104) to obtain a single-cell suspension. About 2000 cells were seeded per 96-well ultra-low-attachment round bottom plate well (Corning, 7007) in Essential 8 medium (Life Technologies, A1517001) supplemented with the ROCK inhibitor Y27632 (10 µM, Selleckchem, S1049),. The plate was then centrifuged at 100g for 3 min and incubated overnight at 37°C with 5% CO2. The next day (day 0) the medium was changed to medium containing DMEM/F12 (Life Technologies, 11320033) with 20% knockout serum (Life Technologies, 10828010), GlutaMax (1:100 Life Technologies 35050079), MEM Non-Essential Amino Acids (1:100 Life Technologies, 11140050), penicillin and streptomycin (1:100, Life Technologies, 15070063), and β-mercaptoethanol supplemented with dorsomorphin (DM; 2.5 µM, Sigma-Aldrich, P5499) and SB-431542 (SB; 10 µM, R&D Systems, 1614) for the first 6 days of differentiation. To induce FP formation, smoothened agonist (SAG; 5 µM, Millipore, 566660), FGF2 (10 ng ml^−1^; R&D Systems), and retinoic acid (RA; Sigma-Aldrich, R2625) were added to the medium starting on day 1, and CHIR 99021 (3 µM; Selleckchem) was added to the media from days 2-6. The medium was changed daily. On days 7 and 8, DM and SB were excluded, and the cultures were be supplemented only with SAG, FGF2, CHIR, and retinoic acid.

### Generation of hSpO from hiPS cells

For the generation of regionalized neural organoids, hiPS cells were incubated with Accutase (Innovative Cell Technologies, AT104) at 37°C for 7 min and dissociated into single cells. For aggregation into organoids, about 3 x 10^6^ single hiPS cells were seeded per AggreWell-800 plate well in Essential 8 medium supplemented with the ROCK inhibitor Y27632 (10 µM, Selleckchem, S1049), centrifuged at 100 g for 3 min, and then incubated at 37°C in 5% CO2. On the next day (day 0), organoids consisting of approximately 10,000 cells were collected and transferred into ultra-low attachment plastic dishes (Corning, 3262) in Essential 6 medium (Thermo Fisher Scientific, A1516401) supplemented with patterning molecules. From days 0-6, organoids were maintained in Essential 6 medium supplemented with DM (2.5 µM, Sigma-Aldrich, P5499) and SB (10 µM, R&D Systems, 1614). On days 5-6, CHIR 99021 (3 µM; Selleckchem) was added. On day 7, organoids were transferred to a neural medium supplemented with EGF (R&D Systems, 236-EG), Retinoic acid (0.1 µM, Sigma-Aldrich, R2625), and CHIR (3 µM, Selleckchem, S1263). On day 22, the media was changed to neural medium with N-2 supplement (Life Technologies, 17502048) supplemented with BDNF (20 ng ml−1; Peprotech, 450-02), cAMP (50 nM; Sigma-Aldrich, D0627), L-Ascorbic Acid (AA, 200 nM; Wako, 321-44823), IGF-1 (10 ng ml^-1^, PeproTech, 100-11) until the end of experiments. From days 22-28, DAPT (2.5 µM; STEMCELL technologies, 72082) was added.

### Generation of hMA from hFpO and hSpO

To generate hMA to study tissue patterning, day 8 hFpO and day 8 hSpO were transferred to a new well containing neural medium supplemented with EGF (R&D Systems, 236-EG), Retinoic acid (0.1 µM, Sigma-Aldrich, R2625), FGF2 (10 ng ml^−1^; R&D Systems), and CHIR (3 µM, Selleckchem, S1263). Half media changes occurred daily and assembloids were collected and processed at days 3 and day 7 after fusion.

To generate hMA to study axon guidance day 8 hFpO and day 22 hSpO were placed in contact with each on a 0.4 µm 6-well tissue culture insert (Corning, 353090). 2mL of neural medium with N-2 supplement (Life Technologies, 17502048) with supplements BDNF (20 ng ml−1; Peprotech, 450-02), cAMP (50 nM; Sigma-Aldrich, D0627), L-Ascorbic Acid (AA, 200 nM; Wako, 321-44823), IGF-1 (10 ng ml^-1^, PeproTech, 100-11), and DAPT (2.5 µM; STEMCELL Technologies, 72082) was added. Half media changes occurred daily and assembloids were collected and processed on days 1, 2, and 3 after fusion.

### Cryopreservation, immunocytochemistry and tissue clearing

Organoids and assembloids were fixed in 4% paraformaldehyde (PFA in PBS, Electron Microscopy Sciences) overnight and subsequently washed three times in PBS. Organoids and assembloids were either prepared for cryosectioning for 2D imaging or directly processed and imaged in 3D.

For 2D preparation, PFA fixed and rinsed organoids were transferred to a solution containing 30% sucrose and left overnight at 4ºC or until the organoids sank in solution. Next, samples were transferred to OCT (Tissue-Tek OCT Compound 4583, Sakura Finetek) and frozen using dry ice. 30 µm thick sections were cut using a cryostat (Leica).

For immunocytochemistry cryosections and free-floating organoids were then blocked for 1 h at room temperature or overnight at 4°C (1% BSA, 0.3% Triton X-100 diluted in PBS). Samples were subsequently incubated overnight at 4°C with primary antibodies in blocking solution. The next day, cryosections were washed with blocking solution and then incubated with Alexa Fluor secondary antibodies (1:1000 dilution in blocking solution) for 1 hr at room temperature or overnight at 4°C. Following washes, actin filaments were visualized with 488 Alexa Fluor Phalloidin (1:200; Invitrogen, A12379). Nuclei were visualized with Hoechst 33258 (Life Technologies) or RedDot2 (Biotium).

For tissue clearing, samples were dehydrated in increasing concentrations of methanol (20%, 50%, 80%, 100%) for 5 min incubation periods and left in pure methanol overnight at 4°C on a shaker. Dehydrated samples were then transferred to vacuum grease wells on a microscope slide. Residual methanol was aspirated and 1:2 benzyl alcohol/benzyl benzoate (BABB) was added and sealed with a glass coverslip. The following antibodies were used for immunostaining: anti-FOXA2 antibody (1:1000 dilution, rabbit, Abcam, ab108422), anti-TUBB3 antibody (1:500 dilution, rabbit, Cell Signaling Technology, MAB1195), anti-NKX2.2 antibody (1:100 dilution, mouse, DSHB, 74.5A5), anti-NKX6.1 antibody (1:100 dilution, mouse, DSHB, F55A10), anti-hROBO3 antibody (1:100 dilution, goat, R&D Systems, AF3076).

Immunostained sections were imaged on a Zeiss Axioscope widefield microscope and 3D samples were imaged using inverted confocal microscopes (Leica). Images were processed and analyzed using Fiji (ver. 2.14.0). ROBO3^+^ axon projection was measured using methods previously described^7-9^ and directionality of pre-crossing axons (100 µm ROI directly adjacent to the assembled hFpO or hSpO) using the Directionality plugin in Fiji (https://imagej.net/plugins/directionality).

### Real-time qPCR

Three organoids were collected in the same tube and considered as one sample. RNA was isolated using the RNeasy Plus Mini kit (Qiagen, 74136). Template cDNA was prepared by reverse transcription using SuperScriptTM III First-Strand Synthesis SuperMix for qRT-PCR (Thermo Fisher Scientific, 11752250). qPCR was performed using SYBRTM Green PCR Master Mix (Thermo Fisher Scientific, 4312704) on a QuantStudio™ 6 Flex Real-Time PCR System (Thermo Fisher Scientific, 4485689). A table with qPCR primer design is included (Supplementary Table 7).

### Single cell RNA seq library preparation and data analysis

Dissociation of organoids was performed as described previously. Four or five randomly selected organoids or assembloids were pooled to obtain single cell suspension and then incubated in 30 U/mL papain enzyme solution (Worthington Biochemical, LS003126) and 0.4% DNase (12,500 U/mL, Worthington Biochemical, LS2007) at 37°C for 45 min. After enzymatic dissociation, organoids were washed with a solution including protease inhibitor and gently triturated to achieve a single cell suspension. Cells were resuspended in 0.04% BSA/PBS (Millipore-Sigma, B6917-25MG) and filtered through a 70 µm Flowmi Cell Strainer (Bel-Art, H13680-0070). The number of cells was counted and loaded onto on a Chromium Single Cell 3′chip (Chromium Next GEM Chip G Single Cell Kit, 10x Genomics, PN-1000127) and cDNA libraries were generated with a Chromium Next GEM Single Cell 3′ GEM, Library & Gel Bead Kit v3.1 (10x Genomics, PN-1000128 and PN-1000269), according to the manufacturer’s instructions. Each library was sequenced using the Illumina NovaSeq S4 2 × 150 bp by Admera Health. UMI counting was performed by the ‘count’ function (--include-introns=TRUE) in Cell Ranger (v7.1.0 and 7.1.0) with Human (GRCh38) 2020-A as reference. Further downstream analyses were performed using the R package Seurat (v4.3.0). Genes on the X or Y chromosome were removed from the count matrix to avoid biases in clustering due to the sex of the hiPS cell lines. Cells with more than 10,000 or less than 2,000 detected genes, with less than 2,000 UMI counts, or cells with mitochondrial content higher than 15% were excluded. Genes that were not expressed in at least three cells were not included in the analysis. Gene expression was normalized using a global-scaling normalization method (normalization method, ‘LogNormalize’; scale factor, 10,000).

The top 2,000, 1000, and 5000 most variable genes were selected for day 22 hSpO, day 8 hFpO, and day 8 hSpO samples, respectively (selection method, ‘vst’) Genes were scaled (mean = 0 and variance = 1 for each gene). Day 22 hSpO samples (n = 3), D11 hFpO-assembled dSpO samples (n = 3),) and day 11 unassembled-hSpO samples (n = 3), and d8 hFpO samples (n = 3) were each separately integrated using ‘FindIntegrationAnchors’ and ‘IntegrateData’ functions with the default parameter. The top 30 principal components were used for clustering (resolution of 1.0), using the ‘FindNeighbors’ and ‘FindClusters’ functions, and for visualization with UMAP. Clusters were grouped based on the expression of known marker genes.

### Mapping organoid single cell RNA-seq data on to reference atlas

Data for label transfer analysis performed in this study was obtained from GEO.^16,35^ Integration, normalization, and clustering were performed on the full dataset (2,000 variable features, 2,000 integration features, 30 dimensions, .5 resolution, and 15% mitochondrial gene cutoff). Neural progenitors were identified and subsetted based on *SOX2, VIM, and TOP2A* expression. The provided annotations and counts by gene matrices were then processed in the same manner as our data before comparison. Label transfer was then carried out with the primary reference datasets using a CCA projection on the first 30 dimensions and using variable features from the integrated assay of our organoid datasets and the RNA assay of the reference dataset.

### Curation of FP enriched membrane and secreted protein list

Secreted proteins from human FP cells were identified by generating conditioned media from 2D FP cultures. hiPS cells were seeded and cultured until 50% confluency in 10 cm plates. The hFpO differentiation protocol was followed and on the final day of differentiation, the cells were washed three times with DMEM/F12 and then cultured in fresh DMEM/F12. After 14 hours, conditioned media was collected and passed through a 0.22 µm filter to remove cellular debris. Next, 1 mL samples were submitted for LC-MS (Tyroma). Proteins were extracted and reduced/alkylated, followed by double digestion with Lys-C/trypsin, and then peptides were desalted. After concentrations were measured, equal amount of each sample was injected into an Ultimate 3000 nanoLC system connected to Q-Exactive HF-X mass spec. The data was analyzed using Proteome Discoverer software with Sequest and Byonic search nodes.

2284 unique proteins were identified in the mass spectrometry date from hFP conditioned media. Only proteins that appeared in all three samples from separate hiPS cell lines were considered, a total of 1876 proteins. FP enriched proteins were identified by overlapping this list with all markers expressed from single cell gene expression data of subsetted human FP cells. When we considered genes that were identified in >30% of FP cells and had a positive log2FC (compared to all other spinal progenitors), this narrowed the data to 415 genes. These genes were then cross referenced with cellular components GO terms: cell surface GO:0009986, cell periphery GO:0071944, extracellular region GO:0005576, extracellular space GO:0005615, external encapsulating structure GO:0030312, secretory vesicle GO:0099503, and membrane GO:0016020. The final list contained 326 membrane and secreted FP-enriched genes, 258 of which were deemed non-essential for cell viability from DepMap Portal. Non-essential genes were considered to have <50% studies deemed essential in CRISPR (DepMap Public 23Q4+Score, Chronos) and RNAi (Achilles+DRIVE+Marcotte, DE-METER2) screens.

### Identification of FP-gene expression divergence in human versus mouse

Gene orthologs were identified by overlapping gene lists of all marker expression from subsetted FP cells in mouse and human primary atlases^16,35^ (Supplementary Table 4) and cross referenced with published curated 1:1 orthologs lists.^35^ Average expression values were log2-transformed, and scatter plots and Pearson’s correlations were calculated to compare human and mouse. Genes were ranked in descending order based on human/mouse expression. Conservation was determined based on previously published methods with human/mouse gene expression <2-fold considered conserved, 2 to 10-fold considered moderately divergent, and >10-fold considered divergent.

### Generation of arrayed CRISPR KO hiPS cell pools

To screen the human-enriched candidate genes, we generated KO hiPS cell pools for the most divergent expressed genes between human and mouse (>10-fold human-enrichment) with the CRISPR–Cas9 system. Three sgRNAs targeting an early exon of each specific gene were designed and synthesized by Synthego to induce one or more fragment deletions (Supplementary Table 5). hiPS cells were dissociated with Accutase, and then 0.5 million cells were mixed with 300 pmol of sgRNAs and 40 pmol of Cas9 protein (Synthego, SpCas9 2NLS Nuclease (1,000 pmol)). Nucleofection was performed using the P3 Primary Cell 4D-NucleofectorTMX Kit L (Lonza, catalogue no. V4XP-3032), a 4D-nucleofector core unit and the X unit (Lonza, program no. CA-137). Cells were then seeded onto vitronectin coated six-well plates in Essential 8 medium supplemented with the ROCK inhibitor Y27632 (10 µM). Essential 8 medium was used for daily medium change. For genotyping, DNA was extracted from hiPS cells using the DNeasy Blood & Tissue Kit (Qiagen, catalogue no. 69506) and PCR was performed with GoTaq G2 Flexi DNA Polymerase (Promega, catalogue no. M7805) with primers to amplify DNA sequence around the sgRNA targeting sites within the gene of interest (Supplementary Table 6). The PCR products were separated by gel electrophoresis and Image Lab (v.6.0.1) was used for gel image acquisition.

### Midline axon guidance screen

Once the arrayed hiPS-KO lines were cultured for 4 days after nucleofection, organoids were generated from each well in round bottom 96-well plates and differentiated to hFpO as described above. The screen was carried out in the 1205-4 hiPS cell background. Quality control of FP induction in hFpO was assessed on days 8 and 11 of differentiation. Three hFpO from each target gene were pooled and underwent RNA isolation as previously described. The gDNA columns used in RNA purification were taken and eluted to collect genomic DNA to assess knockout efficiency. PCR products were generated from each targeted gene as previously described and were analyzed by Sanger sequencing. ICE and KO-score efficiency was determined for each hFpO cell pool collected (https://www.synthego.com/products/bioinformatics/crispr-analysis). RT-qPCR was performed on each candidate to assess *FOXA2* expression. Additionally, FOXA2 expression was assessed by IHC using methods previously described. Isogenic KO screen hits (*GALNT2, NTN1, SPON1*) were generated from the 1205-4 genetic background from an individual clone isolated from hiPS cell pools. Isogenic KO was confirmed via ICE analysis from isolated gDNA and western blot. FP induction efficiency was assessed via RT-qPCR as previously described with KO hiPS cell pools. KO pool hFpO were then assembled with unedited d22 hSpO to form hMA as previously described. The cultures were maintained for 3 days after fusion and fixed and immunostained for FOXA2, ROBO3, and RedDot2 as previously described. Finally, samples were cleared with BABB and imaged with confocal microscope as previously described.

Analysis of the cleared hMA from the screen was carried out in Fiji. hMA images were max-z-projected and oriented so that the hFpO-hSpO boundary was vertical. Plot profiles of ROBO3 fluorescence intensity were generated for 50 µm bins measuring ROBO3 fluorescence intensity within 150 µm pre- and post-hFpO crossing (1.35 µm step size). The data was normalized to the average pre-crossing fluorescence intensity to yield a value signifying the fold-change in ROBO3 fluorescence intensity. Values were min/max normalized using the normalize function in Prism 9. To identify hits from the screen candidates, we performed 2-way ANOVA on the values for 50 µm after the peak fold-change fluorescence intensity of the non-targeting controls. Tukey’s multiple comparison test of the means between non-targeting and targeting sgRNA groups were used to identify statistically significant differences in ROBO3+ axon projection into the hFpO.

### Flow cytometry

To sort GFP^+^ cells from hFpO-assembled or -unassembled hSpO, FACS was conducted on BD-Aria II instruments (Stanford FACS Facility). The ACEA NovoCyte Quanteon 4025 flow cytometer (Stanford FACS Facility) was used to analyze the percentage of GFP+ cells. NovoExpress (v.1.3.0) was used to generate flow cytometry data. Data were analyzed on FlowJo (v.10.8.1).

### Western blotting

Organoids were lysed in Lysis Buffer (50 mM Tris, pH 8.0, 137 mM NaCl, 1 mM EDTA, 1% Triton X-100, and 10% glycerol) supplemented with a Roche complete protease inhibitor set (Roche). After incubation on ice for 20 min and centrifugation at 20,000 × g for 10 minutes, the supernatant was collected for western blot analysis. Conditioned media was mixed with 1/5 volume of 100% Trichloroacetic acid (TCA). After 30 min of incubation on ice, the precipitated proteins were centrifuged at 20,000 x g for 10 min. The pellet was washed with acetone, and resuspended in 1 x SDS Sample Buffer (80 mM Tris, pH 6.8, 2% SDS, 10% Glycerol, 0.0006% Bromophenol blue, 2% β-mercaptoethanol) for western blot analysis. Protein expression was visualized with chemiluminescent substrate (PerkinElmer, NEL113001EA). The following primary antibodies were used: anti-NTN1 antibody (1:1000 dilution, rabbit, Abcam, ab126729), anti-SHH antibody (1:1000 dilution, rabbit, Abcam, ab53281), anti-GAPDH (1:3000 dilution, mouse, Cell Signaling Technology, 2118S), anti-GALNT2 (1:1000 dilution, rabbit, Invitrogen, PA5-21541), and anti-SPON1 (1:1000 dilution, goat, Bio-Techne, AF3135).

### Mouse spinal cord culture

Spinal cords were dissected from E11.5 mouse embryos as previously described^36^ in ice-cold L-15 medium (Gibco, #11415-114), and dorsal regions of the spinal cord were cut into 150 x 150 µm segments, embedded into rattail collagen (Corning, #354236) activated by 12 mM Sodium bicarbonate (Fisher Scientific, #S233-500), and cultured in DMEM/F12 (Gibco, #11-330-032) supplemented with 5% FBS (Gibco, #A52567-01), 1% D-(+)-Glucose (Sigma, #G8769), and Penicillin-Streptomycin-Glutamine (Gibco, #10-378-016), in a 5% CO2, 95% humidity incubator at 37°C for 40 hours.

### Statistics

Data were analyzed with GraphPad Prism (v.9.1.0) unless otherwise indicated. Data are presented as mean ± SEM, unless otherwise indicated. Distribution of the raw data was tested for normality of distribution. Statistical analyses were performed using the Mann–Whitney test and one- or two-way analysis of variance (ANOVA) with multiple comparison tests as indicated. Blinding was performed for imaging experiments and analyses.

### Data availability

The data that support the findings of this study are available on request from the corresponding author. Single cell data is available under Gene Expression Omnibus (GEO) accession number GSE268918 (token: ofkbk-kyidvwtjod). The following public dataset were used: the DepMap Portal (https://depmap.org/portal/). The single-cell RNA sequencing data from primary spinal cells were generated by Rayon et al. (GEO GSE171892).

**Supplementary Figure 1:**
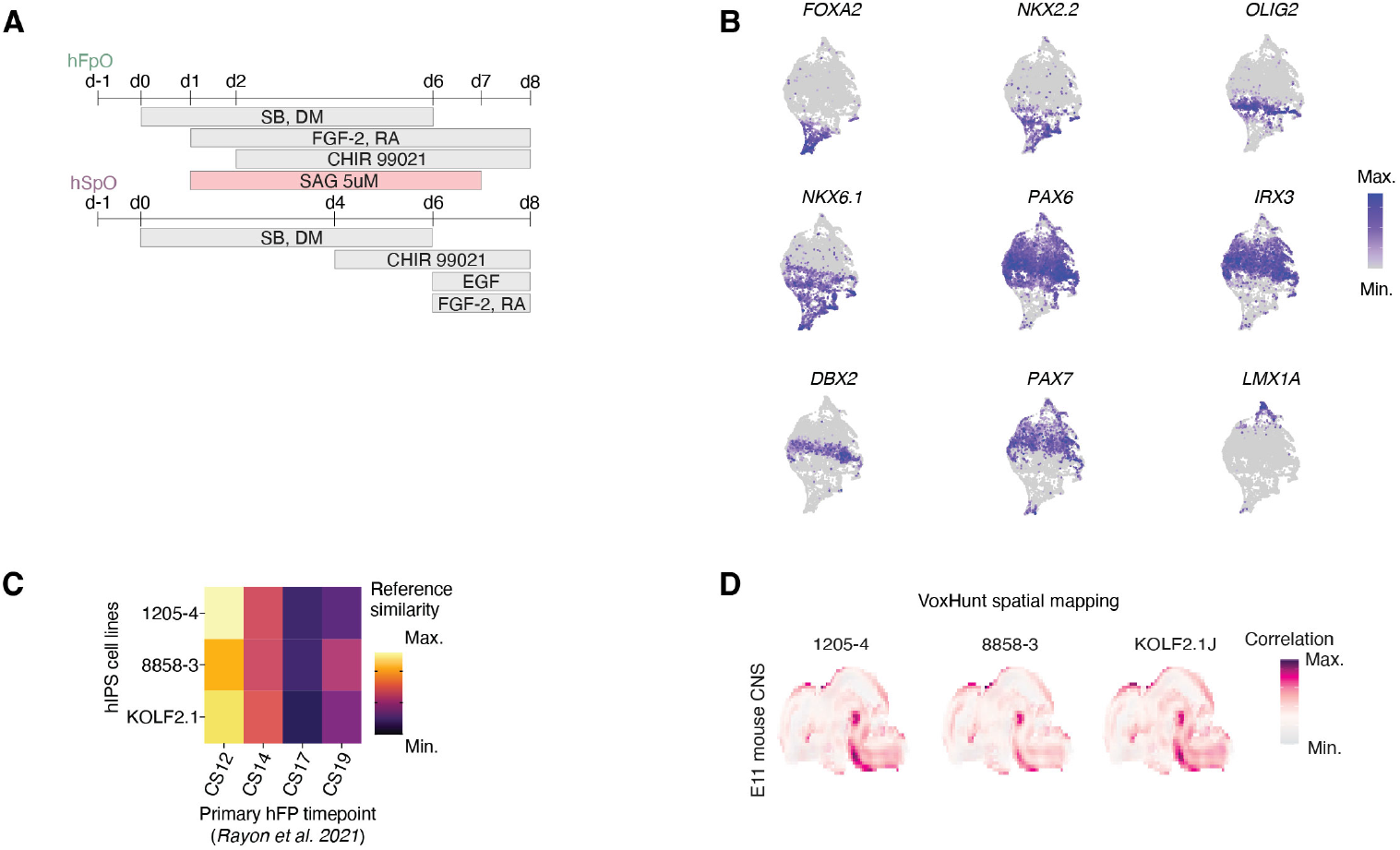
Generation and characterization of 3D hFpO and hSpO. **(A)** Schematic detailing differentiation conditions used for deriving hFpO and hSpO. **(B)** UMAP visualization of dorsal/ventral transcription factor expression in human spinal cord progenitors collected from primary human samples. **(C)** Heatmap displaying the reference similarity of mapped hFpO (n = 3 hiPS cell lines) across timepoints of the primary single cell FP cluster. **(D)** VoxHunt spatial brain mapping of hFpO (n = 3 hiPS cell lines) onto data from E11 mouse nervous system from the Allen Brain Institute.

**Supplementary Figure 2:**
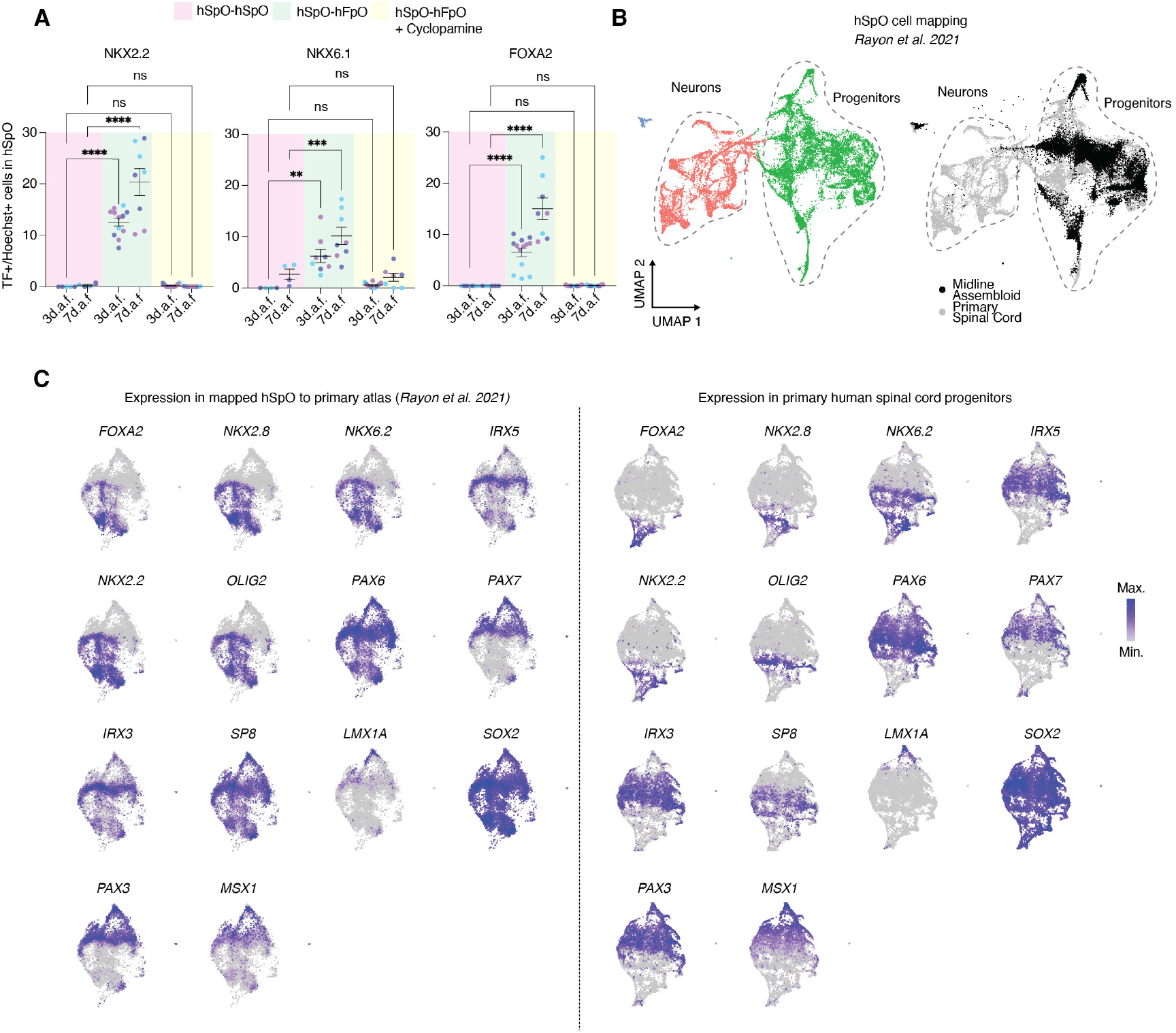
Generation and characterization of hMA for assessing hFP-dependent ventral patterning. **(A)** Quantification of immunocytochemical images for total number of cells positive for ventral transcription factors in hSpO assembled with hSpO (n = 8-13 assembloids, shaded purple), with hFpO (n = 16-21 assembloids, shaded green), and with hFpO in the presence of cyclopamine (CPA, shaded yellow, n = 17-26 assembloids) from 3 separate differentiation experiments (shades of blue). One-way ANOVA with Bonferroni multiple comparison test was used. For NKX2.2; F_5,46_ = 64.23, P < 0.0001, ****P < 0.0001 for hSpO vs hFpO at each timepoint. For NKX6.1, F_5,35_ = 13.41, P < 0.0001, **P = 0.0046 for hSpO vs hFpO at 3 d.a.f and ***P = 0.0006 for hSpO vs hFpO at 7 d.a.f. For FOXA2; F_5,54_ = 49.22, P < 0.0001, ****P < 0.0001 for hSpO vs hFpO at both timepoints. **(B)** UMAP visualization of single cell gene expression of primary human spinal cord neurons and progenitors (left) and mapping of hFpO cells onto the primary atlas (right). Black points indicate the mapped hSpO cells and gray points indicate cells from the primary human spinal cord atlas. **(C)** UMAP visualization of dorsal/ventral transcription factor expression from mapped hSpO (left) compared to human spinal cord progenitors (right). Color scale indicates normalized gene expression level.

**Supplementary Figure 3:**
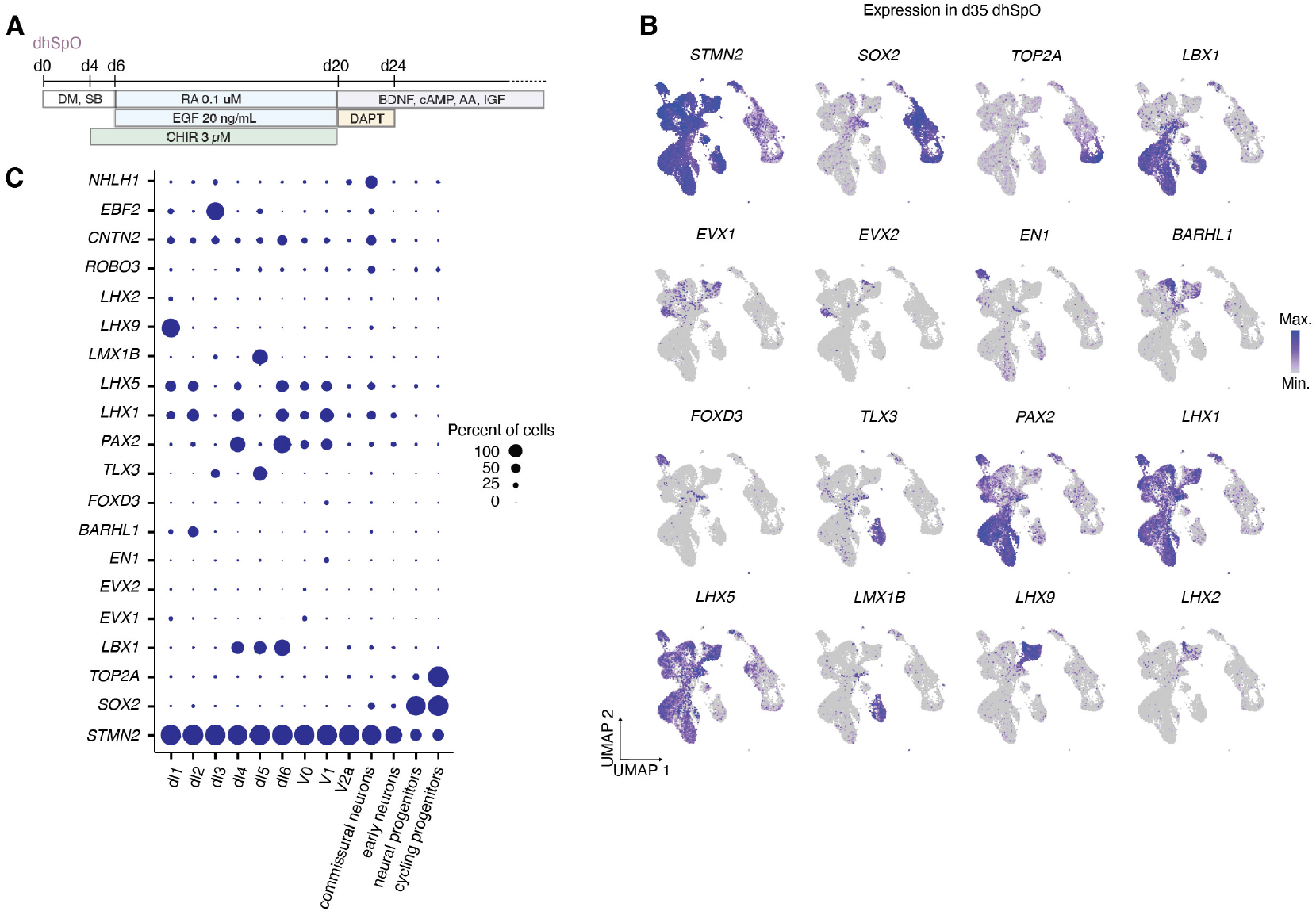
Generation and characterization of hMA to assess hFP-dependent ventral patterning. **(A)** Schematic detailing differentiation conditions used for deriving hSpO. **(B)** UMAP visualization of single cell gene expression of markers enriched in annotated cell populations in day 35 hSpO from 3 hiPS cell lines. **(C)** Dot plots showing the expression of dorsal spinal cord population defining genes. The size of the circle represents the percent of cells expressing each gene.

**Supplementary Figure 4:**
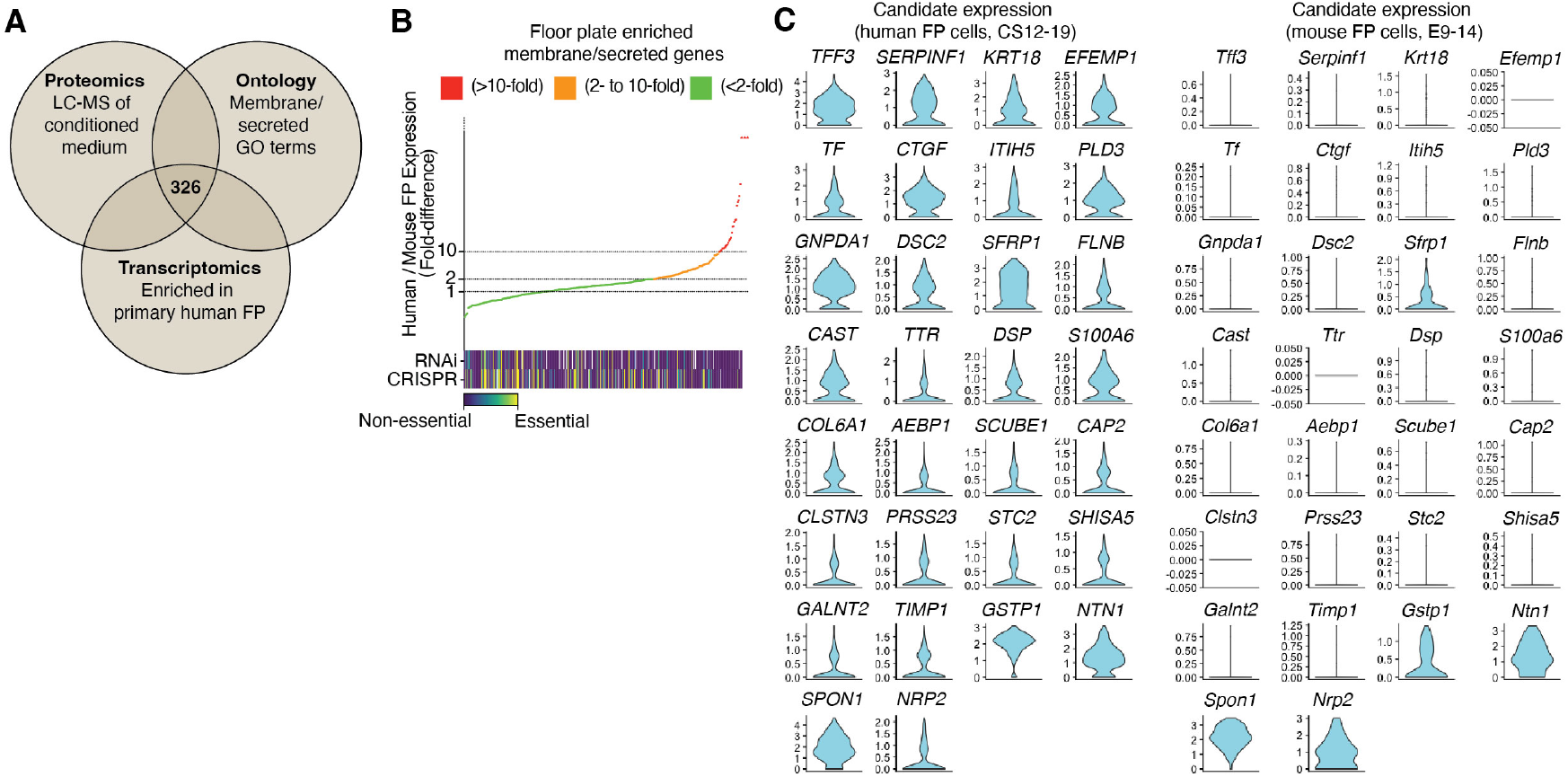
Identification of human enriched FP genes. **(A)** Venn diagram outlining filtering for membrane and secreted genes that are enriched in the FP. The list was curated based on LC-MS identification of proteins from hFpO conditioned media, enriched expression in primary human FP cells from scRNA seq datasets, and presence in membrane/secreted GO terms. **(B)** Graph displaying ranked fold-difference in expression of membrane/secreted FP genes ordered from most conserved to most divergent expression. Heatmap represents the proportion of studies that found the plotted genes to be essential from the DepMap Portal. **(C)** Violin plots of gene expression for the most (>10-fold) divergently expressed membrane/secreted in humans (left) compared to mouse (right) primary FP cells. *SPON1, NTN1, and NRP2* are included as conserved controls between species.

**Supplementary Figure 5:**
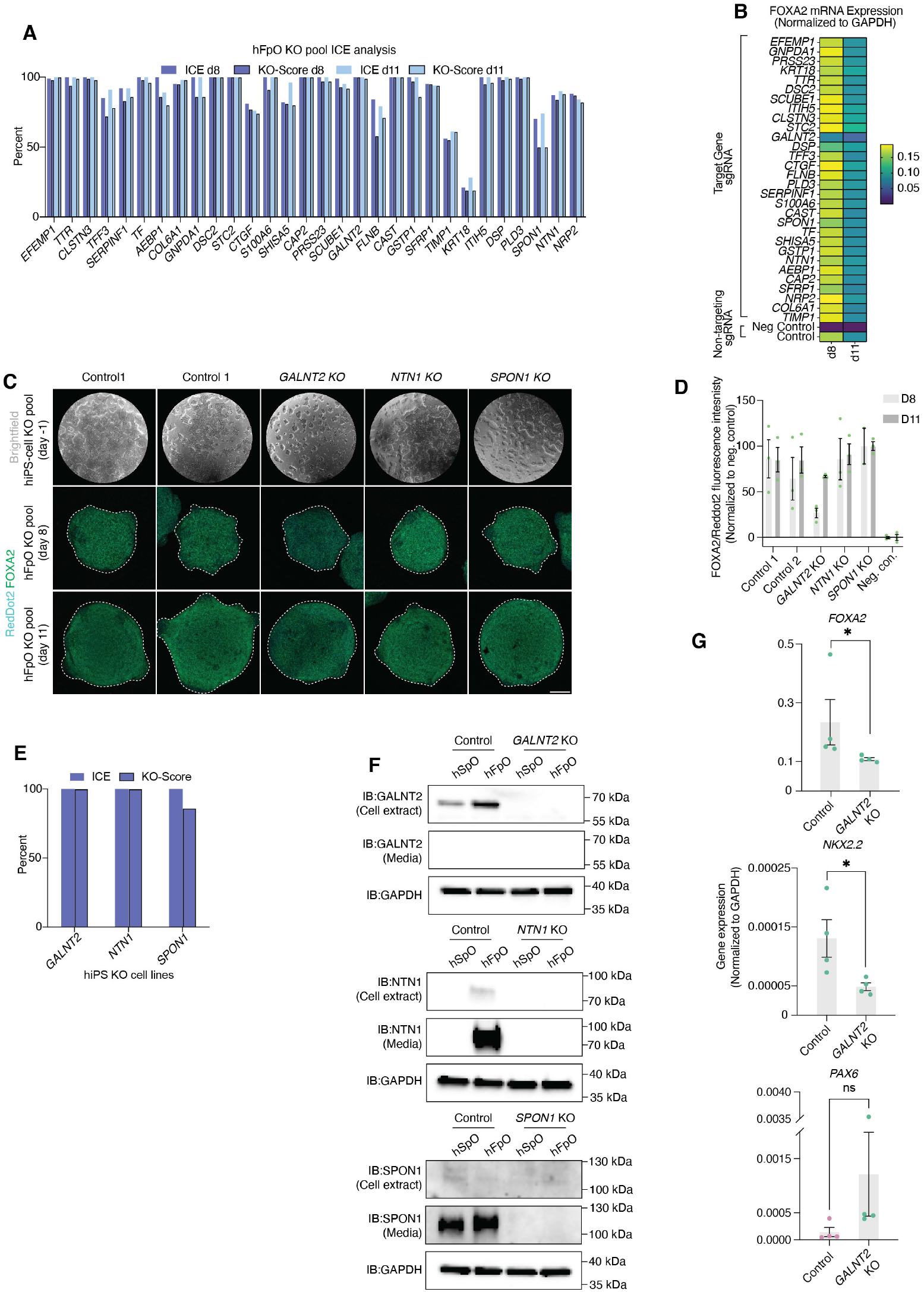
Generation and quality control of pooled and isogenic KO hiPS lines and hFpO. **(A)** Inference of CRISPR edits (ICE; no border) and KO-score (border) of CRISPR edited hFpO at day 8 (dark blue) and day 11 (light blue) of differentiation. **(B)** Gene expression analysis (by RT-qPCR) of *FOXA2* in control and CRISPR edited hFpO at day 8 and 11 of differentiation (pooled mRNA from 3 individual organoids). Color scale indicates the relative expression of *FOXA2* normalized to *GAPDH*. **(C)** Representative brightfield images of control and CRISPR edited hiPS cells (top) and z-projection of immunocytochemical images hFpO (stained for FOXA2 and Reddot2 nuclear stain) at day 8 and 11 of differentiation (bottom). Scale bar 100 μm. **(D)** Quantification of FOXA2/RedDot2 fluorescence intensity of control and CRISPR edited hFpO at day 8 (light gray) and 11 (dark gray) of differentiation (n = 2-3 organoids per condition). Data is normalized to negative control (hSpO). Data is presented as mean ± SEM. **(E)** ICE (no border) and KO-score (border) of isogenic CRISPR edited hiPS cell lines for *GALNT2, NTN1*, and *SPON1* knockout. **(F)** Western blot validation of hSpO and hFpO cell extracts and conditioned media collected from control and isogenic KO for *GALNT2* (top), *NTN1* (middle), and *SPON1* (bottom). GAPDH immunoblotting was used as the sample loading control. **(G)** Gene expression analysis (by RT-qPCR) of *FOXA2, PAX6*, and *NKX2*.*2* in control and isogenic CRISPR edited hFpO at day 8 of differentiation. Each dot represents a separate differentiation experiment normalized to *GAPDH* expression. Data is presented as mean ± SEM. Two-tailed Mann-Whitney test was used. For *FOXA2*: n = 4 control and 4 *GALNT2* KO from 1 hiPS cell line; *P = 0.0286. For *NKX2-2*: n = 4 control and 4 *GALNT2* KO from 1 hiPS cell line; *P = 0.0286. For *PAX6*: n = 4 control and 4 *GALNT2* KO from 1 hiPS cell line; P = 0.0571.

**Supplementary Figure 6:**
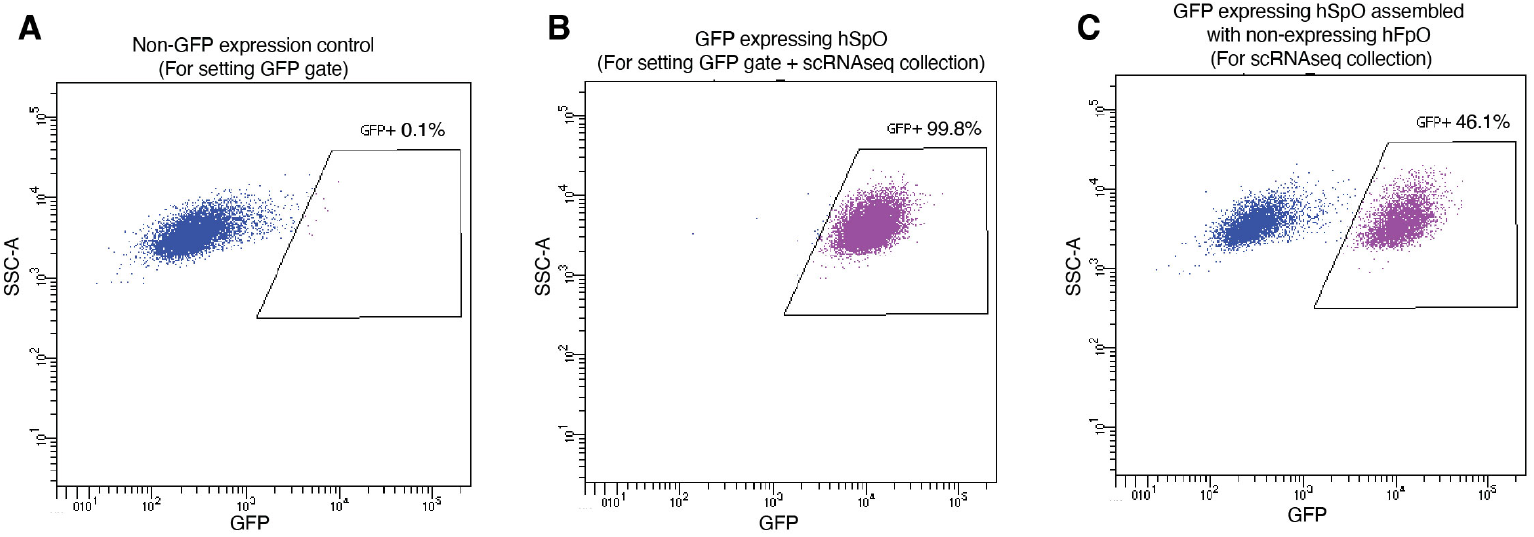
FACS gating strategies for sorting hSpO from hMA. **(A)** Representative FACS plot for negative control (non-GFP expressing hFpO). **(B)** Representative FACS plot for positive control (GFP-expressing hSpO) and setting of gating. **(C)** Representative FACS plot for hMA with hSpO (GFP-expressing) cells sorted from assembled hFpO (non-GFP expressing) cells.

